# Genome-Wide Association Study Meta-Analysis for Parkinson’s Disease Motor Subtypes

**DOI:** 10.1101/2020.06.26.174276

**Authors:** Isabel Alfradique-Dunham, Rami Al-Ouran, Rainer von Coelln, Cornelis Blauwendraat, Emily Hill, Lan Luo, Amanda Stillwell, Emily Young, Manuela Tan, Calwing Liao, Dena G. Hernandez, Lasse Pihlstrom, Donald Grosset, Lisa M. Shulman, Zhandong Liu, Guy A. Rouleau, International Parkinson’s Disease Genomics Consortium (IPDGC), Mike A. Nalls, Andrew B. Singleton, Huw Morris, Joseph Jankovic, Joshua M. Shulman

**Affiliations:** Department of Neurology, Baylor College of Medicine, Houston, TX 77030, USA; Department of Neurology, Washington University, St Louis, MO, 63130; Department of Pediatrics, Baylor College of Medicine, Houston, TX, 77030, USA; Jan and Dan Duncan Neurological Research Institute, Texas Children’s Hospital, Houston, TX, USA; Department of Neurology, University of Maryland School of Medicine, Baltimore, MD, 21201, USA; Molecular Genetics Section, Laboratory of Neurogenetics, National Institute on Aging, National Institutes of Health, Bethesda, MD, 20892, USA; Department of Neurology, Beth Israel Deaconess Medical Center, Harvard Medical School, Boston, MA, 02215, USA; Department of Clinical and Movement Neurosciences, UCL Queen Square Institute of Neurology, University College London, London, UK; UCL Movement Disorders Centre, UCL Queen Square Institute of Neurology, University College London, London, UK; Montreal Neurological Institute, Montréal, Quebec, Canada; Department of Human Genetics, McGill University, Montréal, Quebec, Canada; Department of Neurology, Oslo University Hospital, Oslo, Norway; Department of Neurology, Institute of Neurological Sciences, Queen Elizabeth University Hospital, Glasgow, UK; Department of Neurology and Neurosurgery, McGill University, Montréal, Quebec, Canada; Data Tecnica International, Glen Echo, MD 20812, USA; Parkinson’s Disease Center and Movement Disorders Clinic, Baylor College of Medicine, Houston, TX 77030, USA; Department of Molecular and Human Genetics, Baylor College of Medicine, Houston, TX, 77030, USA; Department of Neuroscience, Baylor College of Medicine, Houston, TX, 77030, USA

**Keywords:** GWAS, Parkinson’s disease, essential tremor, tremor dominant, postural instability/gait difficulty

## Abstract

**OBJECTIVE:** To discover genetic determinants of Parkinson disease (PD) motor subtypes, including Tremor Dominant (TD) and Postural Instability/Gait Difficulty (PIGD) forms.

**METHODS:** In 3,212 PD cases of European ancestry, we performed a genome-wide association study (GWAS) examining two complementary outcome traits derived from the Unified Parkinson’s Disease Rating Scale (UPDRS), including dichotomous motor subtype (TD vs. PIGD) or a continuous tremor / PIGD score ratio. Logistic or linear regression models were adjusted for sex, age of onset, disease duration, and 5 ancestry principal components, followed by meta-analysis.

**RESULTS:** Among 71 established PD risk variants, we detected multiple suggestive associations with PD motor subtype, including *GPNMB* (*rs199347*, p_subtype_ = 0.01, p_ratio_ = 0.03), *SH3GL2* (*rs10756907*, p_subtype_ = 0.02, p_ratio_ = 0.01), *HIP1R* (*rs10847864*, p_subtype_ = 0.02), *RIT2* (*rs12456492*, p_subtype_ = 0.02), and *FBRSL1* (*rs11610045*, p_subtype_ = 0.02). A PD genetic risk score integrating all 71 PD risk variants was also associated with subtype ratio (p = 0.026, ß = −0.04, 95% CI = −0.07, 0). Based on top results of our GWAS, we identify a novel suggestive association at the *STK32B* locus (rs2301857, p_ratio_ = 6.6×10^−7^), which harbors an independent risk allele for essential tremor.

**CONCLUSIONS:** Multiple PD risk alleles may also modify clinical manifestations to influence PD motor subtype. The discovery of a novel variant at *STK32B* suggests a possible overlap between genetic risk for essential tremor and tremor-dominant PD.

## Introduction

Parkinson’s disease (PD) is a clinically heterogeneous disorder.^1-6^ PD subtypes have been described based on common patterns of phenotypic features.^1,7^ One of the earliest and widely used subtyping classifications recognizes tremor dominant (TD) and postural instability/gait difficulty (PIGD) motor subtypes.^8,9^ These subtype categories have implications for disease progression, with prospective studies showing that PIGD is characterized by increased cognitive impairment and decreased response to levodopa.^10,11^ While some studies have sought to identify pathologic correlates for PD motor subtypes^12,13^, the mechanisms underlying these clinical and prognostic differences remain incompletely understood.^7^

A strong genetic contribution to PD etiology is well established, including several rare, monogenic forms of the disease and a large number of common variant PD risk alleles identified in genome-wide association studies (GWAS).^14^ There is mounting evidence for genetic variants as modifiers of PD phenotype as well. Variants in *LRRK2* or *GBA* modify disease motor progression (slower or faster, respectively) and also impact risk of cognitive impairment.^15,16^ Genetic association studies have also nominated genetic modifiers of PD progression, cognitive impairment, age at onset, and risk of insomnia, including established PD risk alleles.^16-25^ Interestingly, *LRRK2(G2019S)* carriers appear to have a higher incidence of the PIGD subtype, despite early reports of asymmetrical tremor as a prominent clinical feature.^15,26^ A recent analysis of 10 PD risk variants from GWAS in a sample of 251 subjects (plus 559 subjects for replication) demonstrated an association of an *SNCA* locus polymorphism with the TD subtype.^20^

We performed a GWAS meta-analysis for PD motor subtype in 3,212 subjects, examining potential associations for 71 established PD risk alleles and further testing for novel modifiers of TD vs. PIGD motor phenotypes.

## Methods

### Participants

All subjects were diagnosed with PD and derived from several North-American and European cohorts (Suppl. Table 1): Baylor College of Medicine (BCM, USA), University of Maryland, Baltimore (UMB, USA) PD Genetics Study, Parkinson’s Progression Markers Initiative (PPMI), Parkinson’s Disease Biomarkers Program (PDBD), Profiling Parkinson’s disease study (PROPARK, Netherlands), Tracking Parkinson’s study (PRoBaND, United Kingdom), and the Oslo Parkinson’s Disease study (Norway). All participants provided written informed consent for genomic studies, including permission for sharing of de-identified data between institutions, prior to enrollment in the respective studies. We obtained all clinical and genetic information with approval of the respective local institutional review boards.

The following data was required for inclusion in this study: sex, age at symptom onset, age at diagnosis, age at first evaluation, and earliest available (baseline) itemized rating using the Unified Parkinson’s Disease Rating Scale (UPDRS) parts 2 and 3 or the equivalent parts of the Movement Disorder Society revised UPDRS version (MDS-UPDRS).^27,28^ Disease duration in years was defined as age at first evaluation minus age at symptom onset. If age of symptom onset was not available, age at diagnosis was used. The BCM cohort included subjects evaluated with either version of the UPDRS, and these subjects were therefore evaluated as separate cohorts (BCM1 and BCM2, see Suppl. Table 1). All other cohorts exclusively used either the UPDRS or the MDS-UPDRS.

### Motor subtypes

PD motor subtypes, TD and PIGD, were determined using previously published algorithms.^1,2^ Subjects are classified as either TD, PIGD or indeterminate using scale-specific cutoffs based on the ratio of tremor score to PIGD score from the UPDRS or MDS-UPDRS parts II and III. Applying these algorithms to our pooled cohort, 383 subjects with a tremor/PIGD score ratio in the indeterminate range could not be assigned to either the TD or PIGD dichotomous trait. As a complementary approach, we therefore used the tremor/PIGD score ratio as a continuous outcome, permitting inclusion of all subjects (including those classified as indeterminate). In order to accommodate subjects with PIGD score = 0 in these analyses, we transformed the tremor/PIGD score ratio as follows:

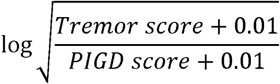

### Genotyping

Genotyping data (all Illumina platform based) were obtained from International Parkinson’s Disease Genomics Consortium (IPDGC) members, collaborators, and public resources. As previously described, all data sets underwent quality control separately, both on individual-level data and variant-level data, as implemented using PLINK v1.90b5.3.^24,29^ Briefly, we excluded individual samples with low or excess heterozygosity or discordant sex. We also excluded ancestry outliers following principal component analysis. We required that SNPs have a minimum call rate of 95%, minor allele frequency (MAF) > 5%, and Hardy-Weinberg equilibrium (HWE) p-values > 1E-04. Imputation was performed using the Michigan imputation server and the Haplotype Reference Consortium (HRC r1.1 2016), with Eagle v2.3 phasing available at: https://imputationserver.sph.umich.edu.

### Statistical analysis

For each study, the imputed genotyped dosages were analyzed using regression, implemented in RVTESTS.^30^ Logistic regression was employed for the dichotomous motor subtype trait (TD vs. PIGD), and linear regression was employed for the continuous tremor/PIGD score ratio trait. Both models were controlled for age of onset, sex, disease duration, and the first 5 ancestry principal components. Fixed effects meta-analysis combining the summary statistics from the 8 studies was performed using METAL with default parameters.^31^ For the GWAS analyses, we computed Lambda_1000_=0.88 for the dichotomous subtype outcome and Lambda=0.99 for the continuous ratio trait. For the candidate analysis of PD-risk alleles, 71 variants had an imputation quality > 0.8 in our dataset and were therefore included in our analyses.^32^ The significance threshold was set at p < 0.0007 based on 71 independent tests using the Bonferroni method (p = 0.05 / 71); we secondarily considered p < 0.05 as evidence of a suggestive association. The 71 PD risk variants were also evaluated in combination using a weighted genetic risk score (GRS), implemented in PLINK.^29,32^ For ease of interpretation, GRS scores were converted to Z scores as previously described.^33^ Association with the 2 subtype outcome traits was tested using the formula:

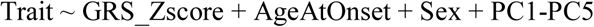

Forest plots and association meta p-values were calculated using the R package metafor.^34^ For the genome-wide analysis, significance was set at p < 5×10^−8^, whereas p < 1×10^−5^ was considered suggestive evidence of association. Locus plots were generated using LocusZoom.^35^ Linkage disequilibrium pruning was performed using the module SNPclip which is part of LDlink application using the default parameters (r^2^ = 0.1 and MAF = 0.01) and a genomic window of 500kb.^36^ For the lookups of variant associations with essential tremor susceptibility, significance was set at p < 0.0013 based on 39 tests. Statistical power was estimated using the Genetic Association Study Power Calculator (http://csg.sph.umich.edu/abecasis/gas_power_calculator/). We performed 2 sets of calculations considering power to detect association of (i) an established PD risk allele (*rs199351*, freq=0.6, risk ratio=1.11) or (ii) a novel variant (*rs10937625*, freq=0.12, risk ratio=1.25). Disease prevalence was set to 0.0041.

### Data Availability Statement

Summary statistics for the analyses presented in this study will be made available on the IPDGC website (http://pdgenetics.org/resources).

## Results

Overall, our study included 3,212 subjects with complete clinical data and genotypes passing all quality control filters (see Methods). Clinical and demographic information along with the frequency of motor subtypes is shown in table 1. The TD subtype was more common than PIGD, but subtype proportions varied between cohorts (table e-1). Consistent with prior reports,^37-39^ the proportion of subjects with TD was inversely related to average disease duration (correlation coefficient −0.57). Due to individuals with indeterminate subtype classification (see Methods), 2,829 subjects were available for the GWAS using the dichotomous subtype trait (TD vs. PIGD), whereas all 3,212 subjects were included in the GWAS for the tremor/PIGD subtype ratio.

**Table 1.** Cohort characteristics.

|  | N (%) or mean (SD) |
| --- | --- |
| N | 3,212 |
| Male | 2,078 (64.7) |
| Age at evaluation (years) | 66.0 (9.5) |
| Age at onset (years) | 61.0 (10.6) |
| Disease duration (years) | 4.1 (4.5) |
| TD subtype | 1,570 (48.9) |
| PIGD subtype | 1,259 (39.2) |
| Indeterminant subtype | 383 (11.9) |

Demographic information and frequency of motor subtypes of the study population combined from all 8 cohorts.

We first examined associations for 71 established PD risk variants with PD motor subtypes. Overall, we identified suggestive associations (p < 0.05) between risk variants at the *GPNMB, SH3GL2, HIP1R, FBRSL1*, and *RIT2* loci and the subtype trait, but none of these associations remained significant following multiple test correction (table 2 and e-2). In 2 out of 5 loci (*GPNMB* and *FBRSL1*), the PD risk-increasing allele was associated with PIGD subtype. Variants at *GPNMB* and *SH3GL2* also showed consistent associations with subtype ratio, but no additional PD risk alleles were associated with this outcome (table e-2). We next integrated genotypes across the 71 PD risk alleles to compute a genetic risk score (GRS) for each subject and examined for association with PD motor subtypes. Indeed, we detected a significant association between the PD GRS and the subtype ratio (p = 0.03, CI= −0.07 - 0.00), although this result appeared to be driven by only 2 out of 8 cohorts included in our meta-analysis (PDBP and BCM2, figure 1). The GRS was not associated with the dichotomous subtype trait (figure e-1).

**Table 2.**
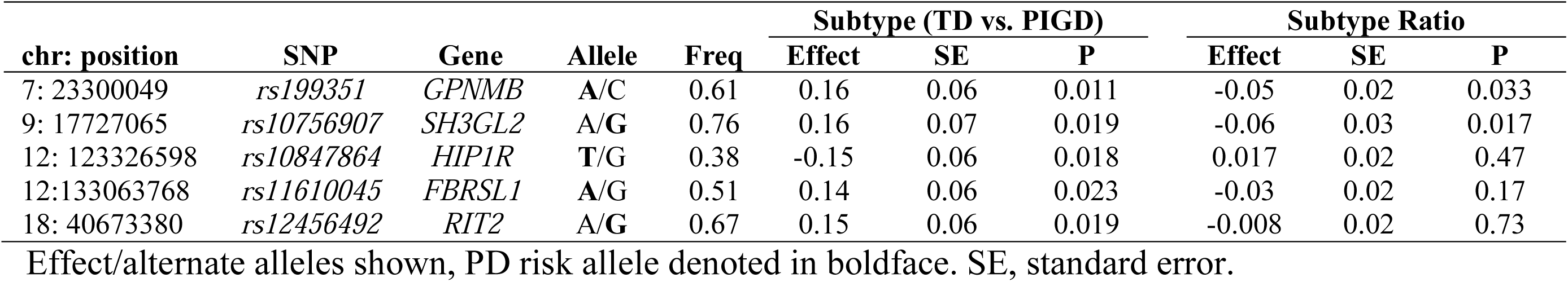
Association of established PD risk variants with PD motor subtype.

**Figure 1.**
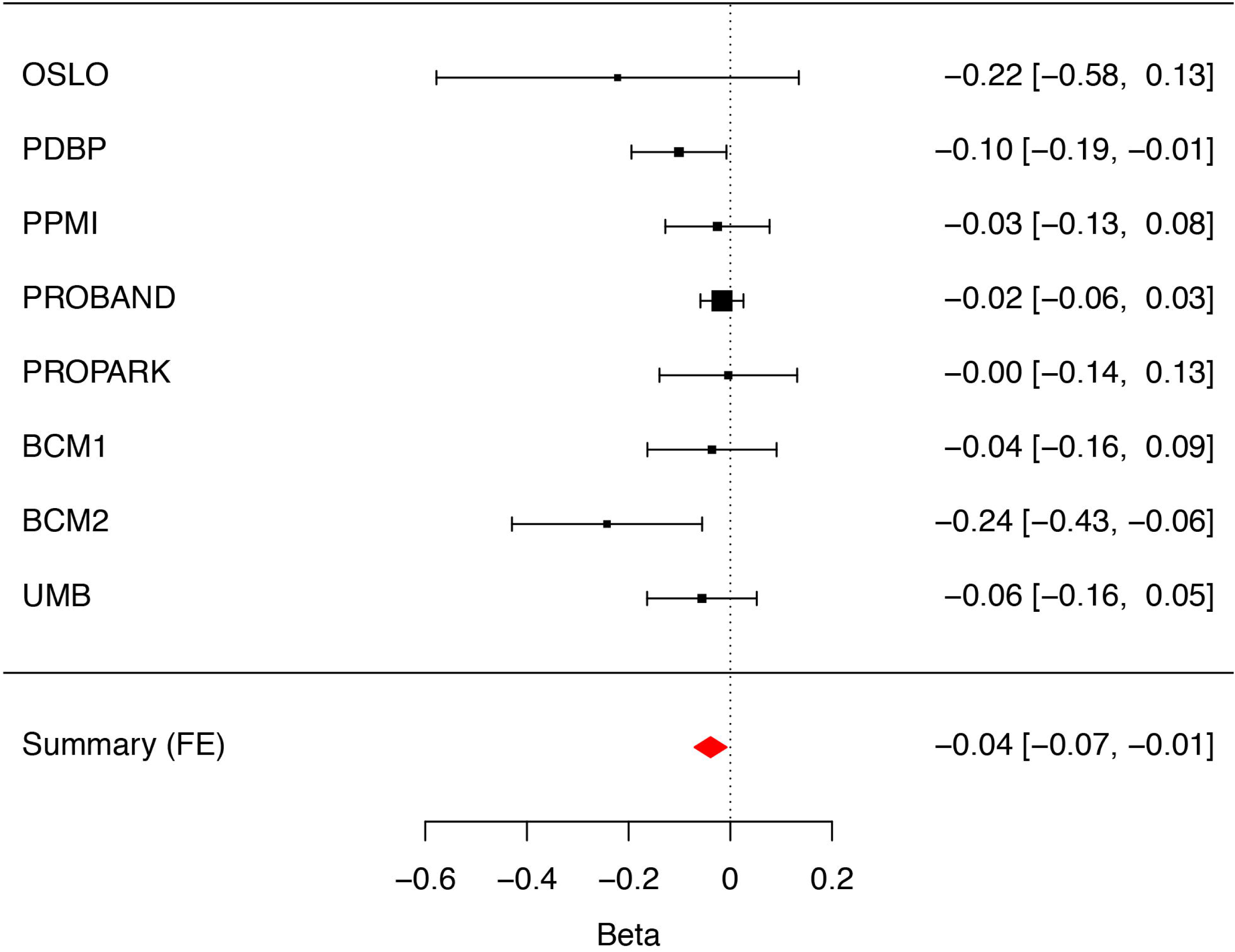
PD genetic risk score associates with tremor/PIGD score ratio. Error bars represent 95% confidence intervals. The size of the black squares represents the effect size from each cohort. The combined estimate for all cohorts is represented by the red diamond with the width of the diamond representing the 95% confidence interval bounds. The summary effect = −0.0389 (p=0.0156).

We next examined the results of our GWAS in order to identify novel candidate modifiers of PD motor subtype. Although no variants reached the genome-wide significance threshold, a number of variants showed suggestive associations (p < 1×10^−5^) with either PD motor subtype or subtype ratio (tables e-3 and e-4). The top variant associated with the subtype ratio outcome is *rs2301857* (p_ratio_ = 6.6×10^−7^), located within an intron of the *STK32B* gene (figure 2). The minor allele, *rs2301857*^*C*^ (freq. = 0.12) was associated with reduced tremor / PIGD score ratio (effect = −0.19). Thus, the minor and major allele for the *rs2301857* SNP are associated with a polarization toward the PIGD vs. TD phenotypes, respectively. In our complementary analysis, the association between *rs2301857* and PD motor subtype was attenuated (p_subtype_ = 0.044). Notably, an association signal at *STK32B* has been previously reported in a GWAS for essential tremor (ET).^40^ Although the lead variant from that study, *rs10937625*, is only 290 kb proximal from the top variant in our analysis, these SNPs do not demonstrate appreciable linkage disequilibrium (R2=0.002, D’=0.184). Based on the common association at the *STK32B* locus, we performed additional analyses to explore for a possible genetic overlap between ET and PD motor subtype. However, neither the *STK32B* variant nor any of the other 5 published ET risk variants^40^ were associated with either of our PD motor subtype traits (table e-5). Lastly, in order to explore for further potential evidence of shared genetic architecture, we reciprocally examined whether any of our top candidate variants (p < 1×10^−5^; n=39 variants) associated with PD motor subtype confers susceptibility for ET, based on lookup of the top results in the largest GWAS completed to date (2807 ET cases / 6441 controls)^40,41^. However, neither *STK32B*^*rs2301857*^ (p=0.18) nor any other top suggestive results from our PD motor subtype GWAS were significantly associated with ET susceptibility.

**Figure 2.**
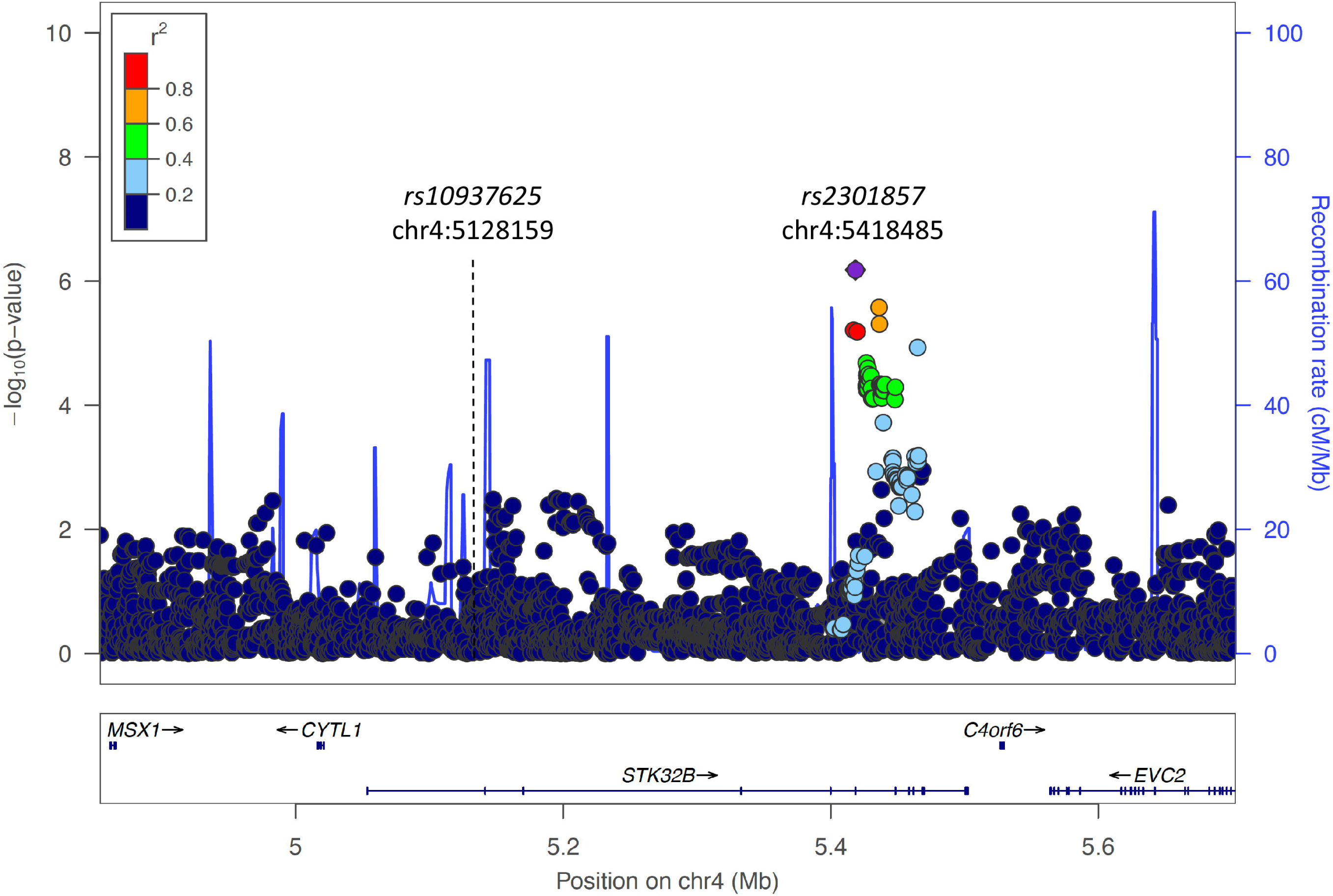
*STK32B* locus association with PD subtype ratio. Locus zoom plot highlighting the association signal at the *STK32B* locus. The top variant associated with PD subtype ratio, *rs2301857*, is highlighted along with other variants in linkage-disequilibrium. Another variant in the same gene, *rs10937625* (dashed line), has been reported as significantly associated with essential tremor (ET), but is not associated with PD subtype ratio. The 2 variants appear independent, and do not show substantial linkage disequilibrium (R2=0.002, D’=0.184).

## Discussion

Identification and characterization of PD subtypes has received increased attention in recent years, with the goal of predicting progression, stratifying patients based on risk of non-motor complications (e.g., dementia), and elucidating mechanisms of disease heterogeneity.^7-9,42^ Recent studies strongly suggest that genetic factors can influence the presence and severity of many varied PD manifestations, and therefore likely influence disease subtypes.^7^ To our knowledge, this is the first GWAS for PD motor subtype. Our results highlight some evidence for 5 established PD risk alleles as potential modifiers of motor subtype, and we further found that a PD GRS including 71 risk variants was associated with subtype ratio. One strength of our analysis was consideration of 2 complementary PD motor subtype outcomes. The TD/PIGD score ratio trait offers a continuous outcome and has the advantage of a larger sample size, since subjects with indeterminate subtype can be considered. On the other hand, by including subjects with a mixed phenotype, it is also possible that the subtype ratio may dilute power to detect the effects of certain variants. In such cases, the dichotomous subtype outcome permits greater contrast between groups of subjects manifesting the TD or PIGD phenotype. In a prior, candidate-based analysis of 10 PD risk alleles in 810 PD cases, a variant at the *SNCA* locus (*rs356182*) was discovered to be associated (p = 0.004, β = 0.7) with a similar TD/PIGD score ratio outcome.^20^ Although we did not replicate that association in our larger sample (n = 3,212, p = 0.18, β=0.03), this may relate to modest differences in the derivation of the subtype score ratio, and additional replication analyses should be undertaken in the future.

While no variants reached genome-wide significance in our GWAS, we identify many loci harboring suggestive associations that may be excellent candidates for follow-up and potential replication. The top result of the subtype ratio GWAS, *rs2301857*, implicates the *STK32B* gene as a possible modifier of PD motor phenotypes. This gene has previously been genetically linked to ET.^40^ The potential relationship between ET and PD has long been a topic of discussion in the field of movement disorders.^43^ While most patients with ET do not develop parkinsonism, at least one study has shown that a prior diagnosis of ET may increase the risk of PD up to 4-fold.^44,45^ A possible genetic link is further suggested by reports of familial co-aggregation of ET and PD.^46^ In another study, PD patients having family members with ET were more likely to exhibit the TD subtype of PD.^47^ Importantly, the variant that we discovered in association with PD motor subtype does not show appreciable linkage-disequilibrium with the previously reported ET susceptibility signal; therefore, these appear to be independent alleles at the *STK32B* gene locus. Thus, while intriguing, our results fall short of providing conclusive evidence of a shared genetic architecture of these two common movement disorders.

Despite including more than 3,000 subjects, statistical power appeared limiting. In fact, we estimate (see Methods) that nearly 14,000 subjects would be required to achieve 80% power to detect a significant association for either a candidate PD risk variant (e.g. *GPNMB*^*rs199351*^) or a novel variant modifier of motor subtype (e.g. *STK32B*^*rs2301857*^). Based on ongoing efforts, we anticipate that sufficiently large cohorts with detailed clinical phenotyping will likely emerge in the next few years. Although the TD and PIGD categories are the earliest and mostly widely used subtype classifications,^8,9^ there are also several notable limitations. The cutoffs employed for differentiating the TD or PIGD subtypes are somewhat arbitrary and without underlying biological or clinical rationale.^1^ In addition, treatment with dopaminergic medication is known to alter the motor UPDRS examination which can thus influence subtype scoring.^39,48^ However, information on medication status and other factors (e.g. dementia) were not universally available for consideration as potential confounders in this analysis. Several studies have also suggested a shift from TD to PIGD subtype along with PD progression, raising questions about the stability of motor subtypes over time.^38,39^ Although our analyses were adjusted for disease duration to account for this in part, future genetic analyses of PD subtypes may benefit from alternative outcome traits that are independent of medication status and disease duration.

## Acknowledgements

We would like to thank all of the subjects who donated their time and biological samples to be a part of this study. We thank J. Marinus and J.J. Van Hilten for use of the PROPARK cohort data and for feedback on the manuscript. The International Parkinson Disease Genomics Consortium (IPDGC, http://pdgenetics.org/partners) is supported in part by the Intramural Research Programs of the National Institute of Neurological Disorders and Stroke (NINDS), the National Institute on Aging (NIA), and the National Institute of Environmental Health Sciences both part of the National Institutes of Health, Department of Health and Human Services; project numbers 1ZIA-NS003154, Z01-AG000949-02 and Z01-ES101986. In addition this work was supported by the Department of Defense (award W81XWH-09-2-0128), and The Michael J Fox Foundation for Parkinson’s Research. This work was supported by National Institutes of Health grants R01NS037167, R01CA141668, P50NS071674, American Parkinson Disease Association (APDA); Barnes Jewish Hospital Foundation; Greater St Louis Chapter of the APDA. The KORA (Cooperative Research in the Region of Augsburg) research platform was started and financed by the Forschungszentrum für Umwelt und Gesundheit, which is funded by the German Federal Ministry of Education, Science, Research, and Technology and by the State of Bavaria. This study was also funded by the German Federal Ministry of Education and Research (BMBF) under the funding code 031A430A, the EU Joint Programme - Neurodegenerative Diseases Research (JPND) project under the aegis of JPND -www.jpnd.eu-through Germany, BMBF, funding code 01ED1406 and iMed – the Helmholtz Initiative on Personalized Medicine. This study is funded by the German National Foundation grant (DFG SH599/6-1) (grant to M.S), Michael J Fox Foundation, and MSA Coalition, USA (to M.S). The French GWAS work was supported by the French National Agency of Research (ANR-08-MNP-012). This study was also funded by France-Parkinson Association, Fondation de France, the French program “Investissements d’avenir” funding (ANR-10-IAIHU-06) and a grant from Assistance Publique-Hôpitaux de Paris (PHRC, AOR-08010) for the French clinical data. This study was also sponsored by the Landspitali University Hospital Research Fund (grant to SSv); Icelandic Research Council (grant to SSv); and European Community Framework Programme 7, People Programme, and IAPP on novel genetic and phenotypic markers of Parkinson’s disease and Essential Tremor (MarkMD), contract number PIAP-GA-2008-230596 MarkMD (to HP and JHu). Institutional research funding IUT20-46 was received of the Estonian Ministry of Education and Research (SK). The McGill study was funded by the Michael J. Fox Foundation and the Canadian Consortium on Neurodegeneration in Aging (CCNA). Mayo Clinic is an American Parkinson Disease Association (APDA) Mayo Clinic Information and Referral Center, an APDA Center for Advanced Research and the Mayo Clinic Lewy Body Dementia Association (LBDA) Research Center of Excellence. OAR is supported by the National Institutes of Health (NIH; R01 NS78086; U54 NS100693, U54 NS110435), the US Department of Defense (W81XWH-17-1-0249), The Little Family Foundation, The Mayo Clinic Functional Genomics of LBD Program the Mayo Clinic Center for Individualized Medicine, and the Michael J. Fox Foundation. Z. K. Wszolek is supported by the NIH, Mayo Clinic Center for Regenerative Medicine, The Sol Goldman Charitable Trust, and Donald G. and Jodi P. Heeringa, the Haworth Family Professorship in Neurodegenerative Diseases fund, and by the Albertson Parkinson’s Research Foundation. This study utilized the high-performance computational capabilities of the Biowulf Linux cluster at the National Institutes of Health, Bethesda, Md. (http://biowulf.nih.gov), and DNA panels, samples, and clinical data from the National Institute of Neurological Disorders and Stroke Human Genetics Resource Center DNA and Cell Line Repository. People who contributed samples are acknowledged in descriptions of every panel on the repository website. We thank the French Parkinson’s Disease Genetics Study Group and the Drug Interaction with genes (DIGPD) study group: Y Agid, M Anheim, F Artaud, A-M Bonnet, C Bonnet, F Bourdain, J-P Brandel, C Brefel-Courbon, M Borg, A Brice, E Broussolle, F Cormier-Dequaire, J-C Corvol, P Damier, B Debilly, B Degos, P Derkinderen, A Destée, A Dürr, F Durif, A Elbaz, D Grabli, A Hartmann, S Klebe, P. Krack, J Kraemmer, S Leder, S Lesage, R Levy, E Lohmann, L Lacomblez, G Mangone, L-L Mariani, A-R Marques, M Martinez, V Mesnage, J Muellner, F Ory-Magne, F Pico, V Planté-Bordeneuve, P Pollak, O Rascol, K Tahiri, F Tison, C Tranchant, E Roze, M Tir, M Vérin, F Viallet, M Vidailhet, A You. We also thank the members of the French 3C Consortium: A Alpérovitch, C Berr, C Tzourio, and P Amouyel for allowing us to use part of the 3C cohort, and D Zelenika for support in generating the genome-wide molecular data. We thank P Tienari (Molecular Neurology Programme, Biomedicum, University of Helsinki), T Peuralinna (Department of Neurology, Helsinki University Central Hospital), L Myllykangas (Folkhalsan Institute of Genetics and Department of Pathology, University of Helsinki), and R Sulkava (Department of Public Health and General Practice Division of Geriatrics, University of Eastern Finland) for the Finnish controls (Vantaa85+ GWAS data). We used genome-wide association data generated by the Wellcome Trust Case-Control Consortium 2 (WTCCC2) from UK patients with Parkinson’s disease and UK control individuals from the 1958 Birth Cohort and National Blood Service. Genotyping of UK replication cases on ImmunoChip was part of the WTCCC2 project, which was funded by the Wellcome Trust (083948/Z/07/Z). UK population control data was made available through WTCCC1. This study was supported by the Medical Research Council and Wellcome Trust disease centre (grant WT089698/Z/09/Z to NW, JHa, and ASc). As with previous IPDGC efforts, this study makes use of data generated by the Wellcome Trust Case-Control Consortium. A full list of the investigators who contributed to the generation of the data is available from www.wtccc.org.uk. Funding for the project was provided by the Wellcome Trust under award 076113, 085475 and 090355. This study was also supported by Parkinson’s UK (grants 8047 and J-0804) and the Medical Research Council (G0700943 and G1100643). Sequencing and genotyping done in McGill University was supported by grants from the Michael J. Fox Foundation, the Canadian Consortium on Neurodegeneration in Aging (CCNA), the Canada First Research Excellence Fund (CFREF), awarded to McGill University for the Healthy Brains for Healthy Lives (HBHL) program and Parkinson’s Society Canada. We thank Jeffrey Barrett and Jason Downing (Illumina Inc) for assistance with the design of the ImmunoChip and NeuroX arrays. DNA extraction work that was done in the UK was undertaken at University College London Hospitals, University College London, who received a proportion of funding from the Department of Health’s National Institute for Health Research Biomedical Research Centres funding. This study was supported in part by the Wellcome Trust/Medical Research Council Joint Call in Neurodegeneration award (WT089698) to the Parkinson’s Disease Consortium (UKPDC), whose members are from the UCL Institute of Neurology, University of Sheffield, and the Medical Research Council Protein Phosphorylation Unit at the University of Dundee. We thank the Quebec Parkinson’s Network (http://rpq-qpn.org) and its members. This work was supported by the Medical Research Council grant MR/N026004/1. The Braineac project was supported by the MRC through the MRC Sudden Death Brain Bank Grant (MR/G0901254) to J.H. P.A.L. was supported by the MRC (grants MR/N026004/1 and MR/L010933/1) and Michael J. Fox Foundation for Parkinson’s Research. Mike A. Nalls’ participation is supported by a consulting contract between Data Tecnica International and the National Institute on Aging, NIH, Bethesda, MD, USA.

## Supplementary Material

Supplementary material includes 5 e-tables, and 1 e-figure.

## Appendix 1: Author Contributions

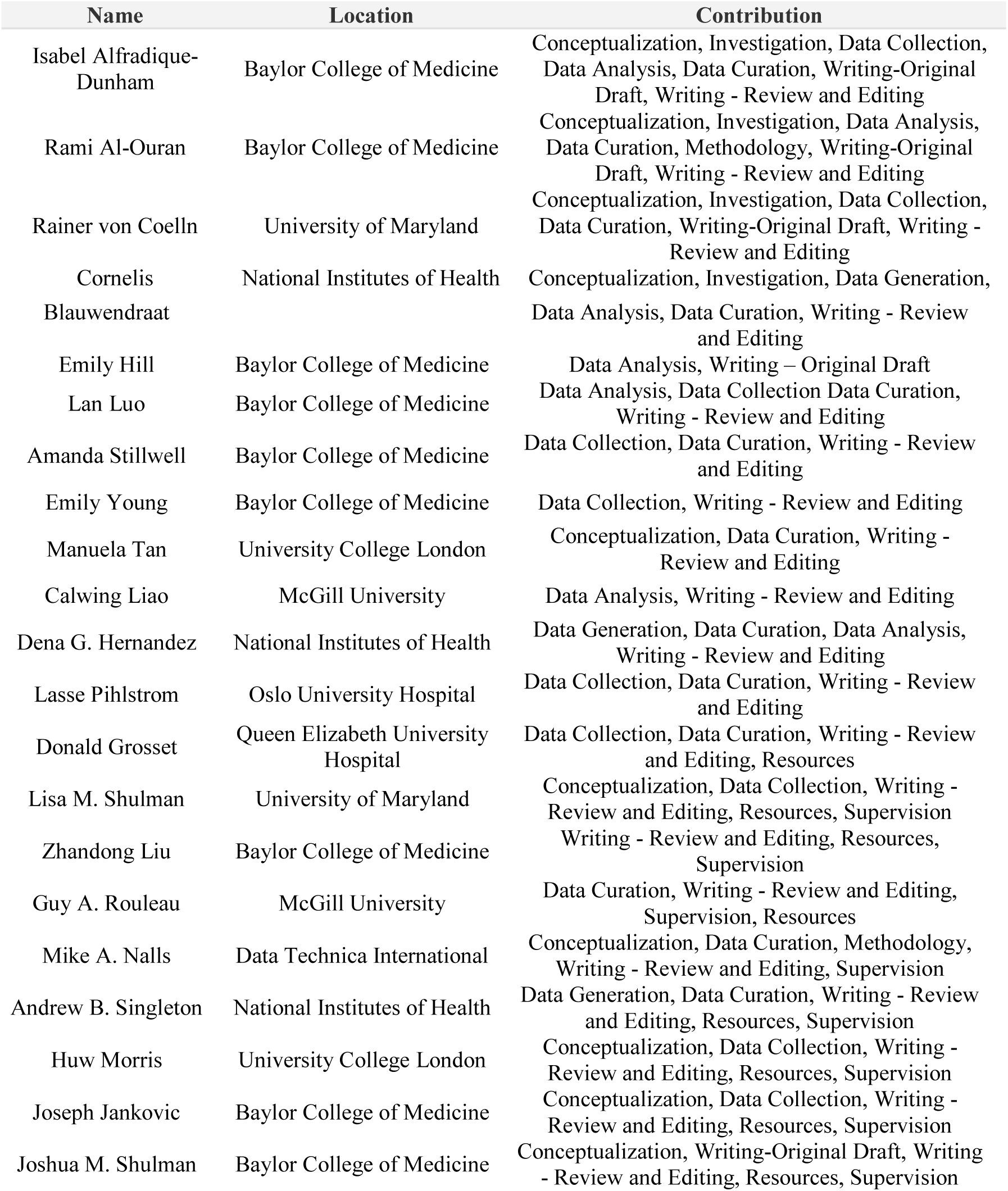

International Parkinson’s Disease Genomics Consortium co-investigators are listed in **Appendix 2**.

## Notes

Relevant Funding Sources: This research was supported in part by the Intramural research Program of the NIH, National institute on Aging and the Parkinson’s UK (Tracking Parkinson’s study J-1101, G-1107, H-1703) and the Medical Research Council (G1100643). GAR receives support from a Canadian Institutes of Health Foundation Grant, and CL is funded by the Vanier Graduate Scholarship. DG receives funding from Parkinson’s UK, the UK National Institute for Health Research, and Glasgow University. JMS was supported by the Huffington Foundation, McGee Foundation, Jan and Dan Duncan Neurological Research Institute at Texas Children’s Hospital, and a Career Award for Medical Scientists from the Burroughs Wellcome Fund. Additional International PD Genomics Consortium funding sources are listed in the Acknowledgements.

### Competing Interest Statement

I. Alfradique-Dunham, R. Al-Ouran, R. von Coelln, C. Blauwendraat, E. Hill, L. Luo, A. Stillwell, E. Young, M. Tan, C. Liao, L. Pihlstrom, J. Marinus, J.J. Van Hilten, D. Grosset, L. Shulman, Z. Liu, G. Rouleau, M. Nalls, A. Singleton, and J. Jankovic report no disclosures relevant to the manuscript. Dr. Nalls consults for Illumina Inc, Lysosomal Therapeutics Inc, the Michael J. Fox Foundation and Vivid Genomics among others. H Morris is employed by UCL and discloses paid consultancy from Biogen, UCB, Abbvie, Denali, Biohaven, Lundbeck; lecture fees/honoraria from Biogen, UCB, C4X Discovery, GE-Healthcare, Wellcome Trust, Movement Disorders Society; Dr Morris is a co-applicant on a patent application related to C9ORF72 - Method for diagnosing a neurodegenerative disease (PCT/GB2012/052140). Dr. Grossett received honoraria from GE Healthcare, Bial Pharma, and Vectura plc, and consultancy fees from The Glasgow Memory Clinic. J. Shulman consults for the Adrienne Helis Malvin & Diana Helis Henry Medical Research Foundations.

