## Supplemental Data for "Genome-Wide Association Study Meta-Analysis for Parkinson’s Disease Motor Subtypes"

**Table e-1. Cohort Clinical Characteristics**

| Cohort | BCM1 | BCM2 | UMB | PPMI | PDBP | PROPARK | PROBAND | OSLO |
|---|---|---|---|---|---|---|---|---|
| Subjects, n | 199 | 83 | 375 | 307 | 429 | 136 | 1641 | 42 |
| Male, n (%) | 124 (62.3%) | 56 (67.5%) | 249 (66.4%) | 210 (68.4%) | 254 (59.2%) | 94 (69.1%) | 1,060 (64.6%) | 31 (73.8%) |
| Age at evaluation, mean (SD) | 60.6 (10.3) | 62.5 (10.1) | 66.7 (9.5) | 63.7 (9.7) | 64.6 (9.3) | 64.9 (7.5) | 67.6 (9.1) | 65.6 (9.2) |
| Age at onset, mean (SD) | 53.6 (9.8) | 56.9 (10.0) | 57.4 (10.5) | 59.5 (9.8) | 57.6 (11.2) | 56.4 (8.9) | 64.5 (9.6) | 57.1 (9.7) |
| Disease duration, mean (SD) | 5.2 (4.7) | 4.4 (3.7) | 4.8 (4.6) | 1.9 (1.8) | 6.9 (7.5) | 8.5 (4.8) | 3.2 (3.0) | 8.5 (6.0) |
| TD subtype, n (%) | 109 (54.8) | 55 (66.3) | 172 (45.9) | 217 (70.7) | 164 (38.2) | 73 (53.7) | 762 (46.4) | 18 (42.9) |
| PIGD subtype, n (%) | 72 (36.2) | 25 (30.1) | 169 (45.1) | 55 (17.9) | 217 (50.6) | 47 (34.6) | 659 (40.2) | 15 (35.7) |
| Subtype ratio, mean (SD) | 0.25 (0.9) | 0.31 (0.9) | 0.15 (1.1) | 0.55 (0.9) | -0.1 (0.9) | 0.22 (0.8) | 0.02 (0.9) | 0.21 (1.2) |

BCM1: Baylor College of Medicine DNA bank with UPDRS; BCM2 = Baylor College of Medicine DNA bank with MDS-UPDRS; UMD = University of Maryland cohort; PPMI = Parkinson's Progression Markers Initiative; PDBP = Parkinson's Disease Biomarkers Program; PROPARK = Profiling Parkinson's disease (Netherlands); PROBAND = Tracking Parkinson's Study (United Kingdom); OSLO = University of Oslo (Norway). PIGD: Postural instability/gait difficulty PD motor subtype, TD: tremor-dominant PD motor subtype.

**Table e-3. Top GWAS results for PD subtype (TD vs. PIGD)**

| chr: position | SNP | Gene | Allele | Freq | Effect | SE | P-value | HetQ | HetP | Direction |
|---|---|---|---|---|---|---|---|---|---|---|
| 1: 47557933 | *rs116504637* | *CYP4Z1* | T/G | 0.74 | 0.35 | 0.07 | 9.83E-07 | 12.9 | 0.33 | +++++++- |
| 5: 161678799 | *rs11949046* | *GABRG2* | T/C | 0.89 | 0.54 | 0.11 | 1.57E-06 | 31 | 0.21 | ++???+++ |
| 16: 19502663 | *rs78926797* | *TMC5* | A/C | 0.60 | -0.3 | 0.06 | 2.08E-06 | 0 | 0.8 | -------- |
| 6: 125107499 | *rs6932127* | *NKAIN2* | T/C | 0.89 | -0.46 | 0.1 | 3.01E-06 | 20.7 | 0.27 | --+-?--- |
| 11: 22583068 | *rs55971529* | *FANCF* | A/G | 0.97 | -1.08 | 0.23 | 3.95E-06 | 51.6 | 0.1 | +????--- |
| 1: 77190938 | *rs988295487* | *ST6GALNAC3* | C/G | 0.96 | -0.84 | 0.18 | 4.98E-06 | 16.8 | 0.31 | --???--- |
| 16: 82279626 | *rs13330839* | *MPHOSPH6* | A/G | 0.87 | -0.47 | 0.1 | 7.00E-06 | 42.1 | 0.12 | +++??--- |
| 6: 134657949 | *rs9483680* | *SGK1* | A/C | 0.9 | 0.49 | 0.11 | 9.69E-06 | 57.3 | 0.04 | +-+??+++ |

Direction of association shown (left to right) for BCM1, BCM2, OSLO, PDBP, PPMI, PROBAND, PROPARK, and UMD.

**Table e-4. Top GWAS results for tremor/PIGD subtype ratio**

| chr: position | SNP | Gene | Allele | Freq | Effect | SE | P-value | HetQ | HetP | Direction |
|---|---|---|---|---|---|---|---|---|---|---|
| 4: 5418485 | *rs2301857* | *STK32B* | C/T | 0.12 | -0.19 | 0.04 | 6.60E-07 | 36 | 0.14 | -+------ |
| 21: 20786475 | *rs79890063* | | C/A | 0.12 | -0.22 | 0.04 | 6.96E-07 | 0 | 0.58 | --+??--- |
| 10: 103048995 | *rs12242050* | *BTRC* | G/T | 0.76 | -0.14 | 0.03 | 8.44E-07 | 13.8 | 0.32 | -++----- |
| 4: 82616045 | *rs6535273* | *RASGEF1B* | C/T | 0.39 | 0.11 | 0.02 | 1.37E-06 | 0 | 0.96 | ++++++++ |
| 13: 43239116 | *rs117046214* | *TNFSF11* | G/A | 0.07 | -0.24 | 0.05 | 1.43E-06 | 60.3 | 0.03 | ---??--- |
| 8: 8387903 | *rs10109703* | *SGK223* | C/A | 0.64 | -0.13 | 0.03 | 1.44E-06 | 16.5 | 0.31 | ---??--- |
| 13: 110127965 | *rs17488950* | *LINC00676* | C/T | 0.07 | -0.27 | 0.06 | 1.46E-06 | 19.5 | 0.29 | -+-??--- |
| 4: 86019236 | *rs76377494* | *WDFY3-AS2* | C/T | 0.07 | -0.22 | 0.05 | 1.89E-06 | 0 | 0.6 | --+-+--- |
| 22: 46008464 | *rs9626407* | *FBLN1* | G/A | 0.04 | -0.35 | 0.07 | 1.97E-06 | 0 | 0.59 | ---??-+- |
| 12: 62440045 | *rs116917617* | *FAM19A2* | C/T | 0.98 | -0.46 | 0.1 | 2.11E-06 | 0 | 0.45 | ---??--- |
| 19: 14878196 | *rs538015403* | *EMR3* | G/A | 0.02 | -0.41 | 0.09 | 2.56E-06 | 40 | 0.11 | -------- |
| 7: 148211886 | *rs2717804* | *C7orf33* | G/A | 0.36 | -0.12 | 0.03 | 2.67E-06 | 0 | 0.45 | ---??--- |
| 4: 163749214 | *rs139714736* | *NAF1* | G/A | 0.03 | 0.41 | 0.09 | 3.01E-06 | 69.5 | 0.006 | +-+??+++ |
| 7: 5794044 | *rs6958272* | *RNF216* | C/T | 0.92 | -0.22 | 0.05 | 3.01E-06 | 0 | 0.58 | +--??--- |
| 16: 15972631 | *rs11644635* | *FOPNL* | C/T | 0.39 | -0.13 | 0.03 | 3.14E-06 | 25.5 | 0.24 | +++??--- |
| 12: 107303612 | *rs7955642* | *RIC8B* | C/T | 0.91 | 0.19 | 0.04 | 5.43E-06 | 54 | 0.03 | ++--++++ |
| 8: 140449676 | *rs117250300* | *KCNK9* | C/T | 0.01 | -0.61 | 0.14 | 5.70E-06 | 56.7 | 0.04 | ---??--- |
| 4: 116121350 | *rs2892663* | *NDST4* | C/T | 0.5 | 0.1 | 0.02 | 5.73E-06 | 32.4 | 0.17 | ++++-+++ |
| 8: 22906180 | *rs7834008* | *TNFRSF10B* | C/T | 0.79 | 0.14 | 0.03 | 5.98E-06 | 26.2 | 0.22 | ++-+++++ |
| 5: 5224692 | *rs2913614* | *ADAMTS16* | T/A | 0.54 | 0.15 | 0.03 | 6.33E-06 | 0 | 0.42 | +++??+++ |
| 2: 47966677 | *rs75342628* | *MSH6* | C/A | 0.93 | 0.22 | 0.05 | 6.38E-06 | 67.2 | 0.009 | -+-??+++ |
| 2: 72300987 | *rs143548133* | *CYP26B1* | C/T | 0.01 | 0.57 | 0.13 | 7.02E-06 | 11.2 | 0.34 | +--??+-+ |
| 6: 134657949 | *rs9483680* | *SGK1* | C/A | 0.9 | -0.19 | 0.04 | 7.02E-06 | 0 | 0.76 | -+-??--- |
| 11: 132873644 | *rs2336592* | *OPCML* | G/C | 0.63 | -0.12 | 0.03 | 7.82E-06 | 0 | 0.61 | ---??--- |
| 1: 105497011 | rs17015314 | none | G/A | 0.88 | -0.17 | 0.04 | 7.90E-06 | 0 | 0.44 | ---??--- |
| 3: 75446661 | rs113942635 | FAM86DP | C/T | 0.01 | 0.81 | 0.18 | 8.00E-06 | 0 | 0.5 | --???+++ |
| 10: 56421400 | rs117238693 | PCDH15 | C/T | 0.98 | 0.43 | 0.1 | 8.04E-06 | 0 | 0.88 | +-++?+++ |
| 11: 103916643 | rs77942836 | PDGFD | G/A | 0.02 | -0.53 | 0.12 | 8.11E-06 | 47 | 0.09 | ---??--+ |
| 2: 57141732 | rs56242312 | none | G/A | 0.9 | -0.19 | 0.04 | 8.18E-06 | 12.3 | 0.34 | -++??--- |
| 19: 45887928 | rs7257687 | PPP1R13L | C/T | 0.03 | -0.30 | 0.07 | 8.30E-06 | 0 | 0.89 | -++----- |
| 13: 87927123 | rs72636490 | MIR4500HG | C/A | 0.08 | -0.21 | 0.05 | 8.59E-06 | 22.6 | 0.26 | ---??--- |
| 14: 47229012 | rs75477320 | RPL10L | C/T | 0.13 | 0.16 | 0.04 | 9.07E-06 | 40.2 | 0.11 | -+++++++ |
| 18: 5197857 | rs80164227 | C18orf42 | G/A | 0.97 | -0.43 | 0.1 | 9.15E-06 | 8.9 | 0.34 | +--??--- |
| 17: 72542970 | rs118076379 | CD300LD | G/A | 0.01 | 0.59 | 0.13 | 9.49E-06 | 0 | 0.69 | +++??+++ |
| 3: 142015590 | rs1552340 | XRN1 | G/A | 0.42 | 0.1 | 0.02 | 9.73E-06 | 41.8 | 0.1 | +++--+++ |
| 7: 132040533 | rs66792854 | PLXNA4 | G/A | 0.06 | 0.25 | 0.06 | 9.96E-06 | 0 | 0.45 | +++??+++ |

Direction of association shown (left to right) for BCM1, BCM2, OSLO, PDBP, PPMI, PROBAND, PROPARK, and UMD.

**Table e-5. ET-risk variant associations with PD motor subtype.**

| | | | | | Subtype (TD vs. PIGD) | | | Subtype Ratio | | |
|---|---|---|---|---|---|---|---|---|---|---|
| **chr: position** | **SNP** | **Gene** | **Allele** | **Freq** | **Effect** | **SE** | **P** | **Effect** | **SE** | **P** |
| 4: 5128159 | rs10937625 | *STK32B* | T/C | 0.75 | 0.02 | 0.07 | 0.79 | -0.01 | 0.03 | 0.78 |
| 4: 24362541 | rs17590046 | *PPARG C1A* | T/C | 0.8 | -0.08 | 0.08 | 0.3 | -0.02 | 0.03 | 0.52 |
| 10: 68845715 | rs12764057 | *CTNNA3* | T/G | 0.62 | 0.06 | 0.07 | 0.35 | -0.04 | 0.03 | 0.14 |
| 10: 68850419 | rs10822974 | *CTNNA3* | A/G | 0.54 | 0.06 | 0.06 | 0.31 | -0.05 | 0.03 | 0.07 |
| 10: 68917164 | rs7903491 | *CTNNA3* | A/G | 0.45 | 0.09 | 0.06 | 0.16 | -0.04 | 0.03 | 0.14 |

**Figure e-1. PD genetic risk score association with motor subtype (TD vs. PIGD).**

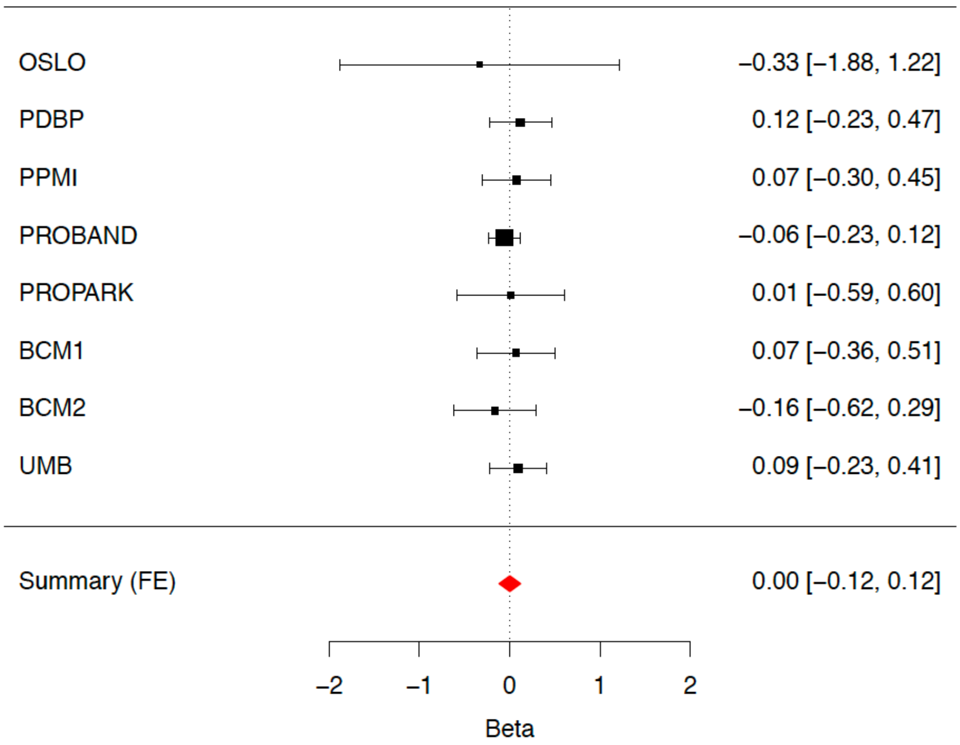


Meta-analysis evaluating for potential association of our genetic risk score and categorically determined PD motor subtypes in all available cohorts. The error bars represent 95% confidence intervals. The size of the black squares represents the effect size from each cohort. The combined estimate for all cohorts is represented by the red diamond with the width of the diamond representing the 95% confidence interval bounds. The summary effect = 0.00157 (p=0.979).
