## Appendix 2 for "Genome-Wide Association Study Meta-Analysis for Parkinson’s Disease Motor Subtypes"

The following IPDGC co-investigators contributed to this project by contributing datasets and reviewing the manuscript:

**United Kingdom:** Alastair J Noyce (Preventive Neurology Unit, Wolfson Institute of Preventive Medicine, QMUL, London, UK and Department of Molecular Neuroscience, UCL, London, UK), Rauan Kaiyrzhanov (Department of Molecular Neuroscience, UCL Institute of Neurology, London, UK), Ben Middlehurst (Institute of Translational Medicine, University of Liverpool, Liverpool, UK), Demis A Kia (UCL Genetics Institute; and Department of Molecular Neuroscience, UCL Institute of Neurology, London, UK), Manuela Tan (Department of Clinical Neuroscience, University College London, London, UK), Henry Houlden (Department of Molecular Neuroscience, UCL Institute of Neurology, London, UK), Catherine Storm (Department of Clinical and Movement Neurosciences, UCL Queen Square Institute of Neurology, London, UK), Huw R Morris (Department of Clinical Neuroscience, University College London, London, UK), Helene Plun-Favreau (Department of Molecular Neuroscience, UCL Institute of Neurology, London, UK), Peter Holmans (Biostatistics & Bioinformatics Unit, Institute of Psychological Medicine and Clinical Neuroscience, MRC Centre for Neuropsychiatric Genetics & Genomics, Cardiff, UK), John Hardy (Department of Molecular Neuroscience, UCL Institute of Neurology, London, UK), Daniah Trabzuni (Department of Molecular Neuroscience, UCL Institute of Neurology, London, UK; Department of Genetics, King Faisal Specialist Hospital and Research Centre, Riyadh, 11211 Saudi Arabia), John Quinn (Institute of Translational Medicine, University of Liverpool, Liverpool, UK), Vivien Bubb (Institute of Translational Medicine, University of Liverpool, Liverpool, UK), Kin Y Mok (Department of Molecular Neuroscience, UCL Institute of Neurology, London, UK), Kerri J. Kinghorn (Institute of Healthy Ageing, Research Department of Genetics, Evolution and Environment, University College London, London, UK), Nicholas W Wood (UCL Genetics Institute; and Department of Molecular Neuroscience, UCL Institute of Neurology, London, UK), Patrick Lewis (University of Reading, Reading, UK), Sebastian R Schreglmann (Department of Molecular Neuroscience, UCL Institute of Neurology, London, UK), Ruth Lovering (University College London, London, UK), Lea R'Bibo (Department of Molecular Neuroscience, UCL Institute of Neurology, London, UK), Claudia Manzoni (University of Reading, Reading, UK), Mie Rizig (Department of Molecular Neuroscience, UCL Institute of Neurology, London, UK), Mina Ryten (Department of Molecular Neuroscience, UCL Institute of Neurology, London, UK), Sebastian Guelfi (Department of Molecular Neuroscience, UCL Institute of Neurology, London, UK), Valentina Escott-Price (MRC Centre for Neuropsychiatric Genetics and Genomics, Cardiff University School of Medicine, Cardiff, UK), Viorica Chelban (Department of Molecular Neuroscience, UCL Institute of Neurology, London, UK), Thomas Foltynie (UCL Institute of Neurology, London, UK), Nigel Williams (MRC Centre for Neuropsychiatric Genetics and Genomics, Cardiff, UK), Karen E. Morrison (Faculty of Medicine, University of Southampton, UK), Carl Clarke (University of Birmingham, Birmingham, UK and Sandwell and West Birmingham Hospitals NHS Trust, Birmingham, UK), Kirsten Harvey (UCL School of Pharmacy, UK), Benjamin M Jacobs (Preventive Neurology Unit, Wolfson Institute of Preventive Medicine, QMUL, London, UK).

**France:** Alexis Brice (Institut du Cerveau et de la Moelle épinière, ICM, Inserm U 1127, CNRS, UMR 7225, Sorbonne Universités, UPMC University Paris 06, UMR S 1127, AP-HP, Pitié-Salpêtrière Hospital, Paris, France), Fabrice Danjou (Institut du Cerveau et de la Moelle épinière, ICM, Inserm U 1127, CNRS, UMR 7225, Sorbonne Universités, UPMC University Paris 06, UMR S 1127, AP-HP, Pitié-Salpêtrière Hospital, Paris, France), Suzanne Lesage (Institut du Cerveau et de la Moelle épinière, ICM, Inserm U 1127, CNRS, UMR 7225, Sorbonne Universités, UPMC University Paris 06, UMR S 1127, AP-HP, Pitié-Salpêtrière Hospital, Paris, France), Jean-Christophe Corvol (Institut du Cerveau et de la Moelle épinière, ICM, Inserm U 1127, CNRS, UMR 7225, Sorbonne Universités, UPMC University Paris 06, UMR S 1127, Centre d'Investigation Clinique Pitié Neurosciences CIC-1422, AP-HP, Pitié-Salpêtrière Hospital, Paris, France), Maria Martinez (INSERM UMR 1220; and Paul Sabatier University, Toulouse, France),

**Germany:** Claudia Schulte (Department for Neurodegenerative Diseases, Hertie Institute for Clinical Brain Research, University of Tübingen, and DZNE, German Center for Neurodegenerative Diseases, Tübingen, Germany), Kathrin

Brockmann (Department for Neurodegenerative Diseases, Hertie Institute for Clinical Brain Research, University of Tübingen, and DZNE, German Center for Neurodegenerative Diseases, Tübingen, Germany), Javier Simón-Sánchez (Department for Neurodegenerative Diseases, Hertie Institute for Clinical Brain Research, University of Tübingen, and DZNE, German Center for Neurodegenerative Diseases, Tübingen, Germany), Peter Heutink (DZNE, German Center for Neurodegenerative Diseases and Department for Neurodegenerative Diseases, Hertie Institute for Clinical Brain Research, University of Tübingen, Tübingen, Germany), Patrizia Rizzu (DZNE, German Center for Neurodegenerative Diseases), Manu Sharma (Centre for Genetic Epidemiology, Institute for Clinical Epidemiology and Applied Biometry, University of Tubingen, Germany), Thomas Gasser (Department for Neurodegenerative Diseases, Hertie Institute for Clinical Brain Research, and DZNE, German Center for Neurodegenerative Diseases, Tübingen, Germany), Susanne A. Schneider (Department of Neurology, Ludwig-Maximilians-University Munich, München, Germany)

**United States of America:** Mark R Cookson (Laboratory of Neurogenetics, National Institute on Aging, Bethesda, USA), Sara Bandres-Ciga (Laboratory of Neurogenetics, National Institute on Aging, Bethesda, MD, USA), Cornelis Blauwendraat (Laboratory of Neurogenetics, National Institute on Aging, Bethesda, MD, USA), David W. Craig (Department of Translational Genomics, Keck School of Medicine, University of Southern California, Los Angeles, USA), Kimberley Billingsley (Laboratory of Neurogenetics, National Institute on Aging, Bethesda, MD, USA), Mary B. Makarious (Laboratory of Neurogenetics, National Institute on Aging, Bethesda, MD, USA), Derek Narendra (Inherited Movement Disorders Unit, National pInstitute of Neurological Disorders and Stroke, Bethesda, MD, USA), Faraz Faghri (Laboratory of Neurogenetics, National Institute on Aging, Bethesda, USA; Department of Computer Science, University of Illinois at Urbana-Champaign, Urbana, IL, USA), J Raphael Gibbs (Laboratory of Neurogenetics, National Institute on Aging, National Institutes of Health, Bethesda, MD, USA), Dena G. Hernandez (Laboratory of Neurogenetics, National Institute on Aging, Bethesda, MD, USA), Kendall Van Keuren-Jensen (Neurogenomics Division, TGen, Phoenix, AZ USA), Joshua M. Shulman (Departments of Neurology, Neuroscience, and Molecular & Human Genetics, Baylor College of Medicine, Houston, Texas, USA; Jan and Dan Duncan Neurological Research Institute, Texas Children's Hospital, Houston, Texas, USA), Hirotaka Iwaki (Laboratory of Neurogenetics, National Institute on Aging, Bethesda, MD, USA), Hampton L. Leonard (Laboratory of Neurogenetics, National Institute on Aging, Bethesda, MD, USA), Mike A. Nalls (Laboratory of Neurogenetics, National Institute on Aging, Bethesda, USA; CEO/Consultant Data Tecnica International, Glen Echo, MD, USA), Laurie Robak (Baylor College of Medicine, Houston, Texas, USA), Jose Bras (Center for Neurodegenerative Science, Van Andel Research Institute, Grand Rapids, Michigan, USA), Rita Guerreiro (Center for Neurodegenerative Science, Van Andel Research Institute, Grand Rapids, Michigan, USA), Steven Lubbe (Ken and Ruth Davee Department of Neurology, Northwestern University Feinberg School of Medicine, Chicago, IL, USA), Steven Finkbeiner (Departments of Neurology and Physiology, University of California, San Francisco; Gladstone Institute of Neurological Disease; Taube/Koret Center for Neurodegenerative Disease Research, San Francisco, CA, USA), Niccolo E. Mencacci (Northwestern University Feinberg School of Medicine, Chicago, IL, USA), Codrin Lungu (National Institutes of Health Division of Clinical Research, NINDS, National Institutes of Health, Bethesda, MD, USA), Andrew B Singleton (Laboratory of Neurogenetics, National Institute on Aging, Bethesda, MD, USA), Sonja W. Scholz (Neurodegenerative Diseases Research Unit, National Institute of Neurological Disorders and Stroke, Bethesda, MD, USA), Xylena Reed (Laboratory of Neurogenetics, National Institute on Aging, Bethesda, MD, USA). Roy N. Alcalay (Department of Neurology, College of Physicians and Surgeons, Columbia University Medical Center, New York, NY, USA, Taub Institute for Research on Alzheimer's Disease and the Aging Brain, College of Physicians and Surgeons, Columbia University Medical Center, New York, NY, USA). Zbigniew K. Wszolek (Department of Neurology, Mayo Clinic Jacksonville, FL, USA), Ryan J. Uitti (Department of Neurology, Mayo Clinic Jacksonville, FL, USA), Owen A. Ross (Departments of Neuroscience & Clinical Genomics, Mayo Clinic Jacksonville, FL, USA).

**Canada:** Ziv Gan-Or (Montreal Neurological Institute and Hospital, Department of Neurology & Neurosurgery, Department of Human Genetics, McGill University, Montréal, QC, H3A 0G4, Canada), Guy A. Rouleau (Montreal Neurological Institute and Hospital, Department of Neurology & Neurosurgery, Department of Human Genetics,

McGill University, Montréal, QC, H3A 0G4, Canada), Lynne Krohn (Montreal Neurological Institute and Hospital, Department of Neurology & Neurosurgery, Department of Human Genetics, McGill University, Montréal, QC, H3A 0G4, Canada), Kheireddin Mufti (Montreal Neurological Institute and Hospital, Department of Neurology & Neurosurgery, Department of Human Genetics, McGill University, Montréal, QC, H3A 0G4, Canada),

**The Netherlands:** Jacobus J van Hilten (Department of Neurology, Leiden University Medical Center, Leiden, Netherlands), Johan Marinus (Department of Neurology, Leiden University Medical Center, Leiden, Netherlands)

**Spain:** Astrid D. Adarmes-Gómez (Instituto de Biomedicina de Sevilla (IBiS), Hospital Universitario Virgen del Rocío/CSIC/Universidad de Sevilla, Seville), Miquel Aguilar (Fundació Docència i Recerca Mútua de Terrassa and Movement Disorders Unit, Department of Neurology, University Hospital Mutua de Terrassa, Terrassa, Barcelona.), Ignacio Alvarez (Fundació Docència i Recerca Mútua de Terrassa and Movement Disorders Unit, Department of Neurology, University Hospital Mutua de Terrassa, Terrassa, Barcelona.),Victoria Alvarez (Hospital Universitario Central de Asturias, Oviedo), Francisco Javier Barrero (Hospital Universitario San Cecilio de Granada, Universidad de Granada), Jesús Alberto Bergareche Yarza (Instituto de Investigación Sanitaria Biodonostia, San Sebastián), Inmaculada Bernal-Bernal (Instituto de Biomedicina de Sevilla (IBiS), Hospital Universitario Virgen del Rocío/CSIC/Universidad de Sevilla, Seville), Marta Blazquez (Hospital Universitario Central de Asturias, Oviedo), Marta Bonilla-Toribio (Instituto de Biomedicina de Sevilla (IBiS), Hospital Universitario Virgen del Rocío/CSIC/Universidad de Sevilla, Seville), Juan A. Botía (Universidad de Murcia, Murcia), María Teresa Boungiorno (Fundació Docència i Recerca Mútua de Terrassa and Movement Disorders Unit, Department of Neurology, University Hospital Mutua de Terrassa, Terrassa, Barcelona.) Dolores Buiza-Rueda (Instituto de Biomedicina de Sevilla (IBiS), Hospital Universitario Virgen del Rocío/CSIC/Universidad de Sevilla, Seville), Ana Cámara (Hospital Clinic de Barcelona), Fátima Carrillo (Instituto de Biomedicina de Sevilla (IBiS), Hospital Universitario Virgen del Rocío/CSIC/Universidad de Sevilla, Seville), Mario Carrión-Claro (Instituto de Biomedicina de Sevilla (IBiS), Hospital Universitario Virgen del Rocío/CSIC/Universidad de Sevilla, Seville), Debora Cerdan (Hospital General de Segovia, Segovia), Jordi Clarimón (Memory Unit, Department of Neurology, IIB Sant Pau, Hospital de la Santa Creu i Sant Pau, Universitat Autònoma de Barcelona and Centro de Investigación Biomédica en Red en Enfermedades Neurodegenerativas (CIBERNED), Madrid),Yaroslau Compta (Hospital Clinic de Barcelona), Monica Diez-Fairen (Fundació Docència i Recerca Mútua de Terrassa and Movement Disorders Unit, Department of Neurology, University Hospital Mutua de Terrassa, Terrassa, Barcelona.), Oriol Dols-Icardo (Memory Unit, Department of Neurology, IIB Sant Pau, Hospital de la Santa Creu i Sant Pau, Universitat Autònoma de Barcelona, Barcelona, and Centro de Investigación Biomédica en Red en Enfermedades Neurodegenerativas (CIBERNED), Madrid), Jacinto Duarte (Hospital General de Segovia, Segovia), Raquel Duran (Centro de Investigacion Biomedica, Universidad de Granada, Granada), Francisco Escamilla-Sevilla (Hospital Universitario Virgen de las Nieves, Instituto de Investigación Biosanitaria de Granada, Granada), Mario Ezquerra (Hospital Clinic de Barcelona), Cici Feliz (Departmento de Neurologia, Instituto de Investigación Sanitaria Fundación Jiménez Díaz, Madrid, Spain), Manel Fernández (Hospital Clinic de Barcelona), Rubén Fernández-Santiago (Hospital Clinic de Barcelona), Ciara Garcia (Hospital Universitario Central de Asturias, Oviedo), Pedro García-Ruiz (Instituto de Investigación Sanitaria Fundación Jiménez Díaz, Madrid), Pilar Gómez-Garre (Instituto de Biomedicina de Sevilla (IBiS), Hospital Universitario Virgen del Rocío/CSIC/Universidad de Sevilla, Seville), Maria Jose Gomez Heredia (Hospital Universitario Virgen de la Victoria, Malaga), Isabel Gonzalez-Aramburu (Hospital Universitario Marqués de Valdecilla-IDIVAL, Santander) ,Ana Gorostidi Pagola (Instituto de Investigación Sanitaria Biodonostia, San Sebastián), Janet Hoenicka (Institut de Recerca Sant Joan de Déu, Barcelona), Jon Infante (Hospital Universitario Marqués de Valdecilla-IDIVAL and University of Cantabria, Santander, and Centro de Investigación Biomédica en Red en Enfermedades Neurodegenerativas (CIBERNED), Silvia Jesús (Instituto de Biomedicina de Sevilla (IBiS), Hospital Universitario Virgen del Rocío/CSIC/Universidad de Sevilla, Seville), Adriano Jimenez-Escrig (Hospital Universitario Ramón y Cajal, Madrid), Jaime Kulisevsky (Movement Disorders Unit, Department of Neurology, IIB Sant Pau, Hospital de la Santa Creu i Sant Pau, Universitat Autònoma de Barcelona, Barcelona, and Centro de Investigación Biomédica en Red en Enfermedades Neurodegenerativas (CIBERNED)), Miguel A. Labrador-Espinosa

(Instituto de Biomedicina de Sevilla (IBiS), Hospital Universitario Virgen del Rocío/CSIC/Universidad de Sevilla, Seville), Jose Luis Lopez-Sendon (Hospital Universitario Ramón y Cajal, Madrid), Adolfo López de Munain Arregui (Instituto de Investigación Sanitaria Biodonostia, San Sebastián), Daniel Macias (Instituto de Biomedicina de Sevilla (IBiS), Hospital Universitario Virgen del Rocío/CSIC/Universidad de Sevilla, Seville), Irene Martínez Torres (Department of Neurology, Instituto de Investigación Sanitaria La Fe, Hospital Universitario y Politécnico La Fe, Valencia), Juan Marín (Movement Disorders Unit, Department of Neurology, IIB Sant Pau, Hospital de la Santa Creu i Sant Pau, Universitat Autònoma de Barcelona, Barcelona, and Centro de Investigación Biomédica en Red en Enfermedades Neurodegenerativas (CIBERNED)), Maria Jose Marti (Hospital Clinic Barcelona), Juan Carlos Martínez-Castrillo (Instituto Ramón y Cajal de Investigación Sanitaria, Hospital Universitario Ramón y Cajal, Madrid), Carlota Méndez-del-Barrio (Instituto de Biomedicina de Sevilla (IBiS), Hospital Universitario Virgen del Rocío/CSIC/Universidad de Sevilla, Seville), Manuel Menéndez González (Hospital Universitario Central de Asturias, Oviedo), Marina Mata (Department of Neurology, Hospital Universitario Infanta Sofía, Madrid, Spain) Adolfo Mínguez (Hospital Universitario Virgen de las Nieves, Granada, Instituto de Investigación Biosanitaria de Granada), Pablo Mir (Instituto de Biomedicina de Sevilla (IBiS), Hospital Universitario Virgen del Rocío/CSIC/Universidad de Sevilla, Seville), Elisabet Mondragon Rezola (Instituto de Investigación Sanitaria Biodonostia, San Sebastián), Esteban Muñoz (Hospital Clinic Barcelona), Javier Pagonabarraga (Movement Disorders Unit, Department of Neurology, IIB Sant Pau, Hospital de la Santa Creu i Sant Pau, Universitat Autònoma de Barcelona, Barcelona, and Centro de Investigación Biomédica en Red en Enfermedades Neurodegenerativas (CIBERNED)), Pau Pastor (Fundació Docència i Recerca Mútua de Terrassa and Movement Disorders Unit, Department of Neurology, University Hospital Mutua de Terrassa, Terrassa, Barcelona.), Francisco Perez Errazquin (Hospital Universitario Virgen de la Victoria, Malaga), Teresa Periñán-Tocino (Instituto de Biomedicina de Sevilla (IBiS), Hospital Universitario Virgen del Rocío/CSIC/Universidad de Sevilla, Seville), Javier Ruiz-Martínez (Hospital Universitario Donostia, Instituto de Investigación Sanitaria Biodonostia, San Sebastián), Clara Ruz (Centro de Investigacion Biomedica, Universidad de Granada, Granada), Antonio Sanchez Rodriguez (Hospital Universitario Marqués de Valdecilla-IDIVAL, Santander), María Sierra (Hospital Universitario Marqués de Valdecilla-IDIVAL, Santander), Esther Suarez-Sanmartin (Hospital Universitario Central de Asturias, Oviedo), Cesar Tabernero (Hospital General de Segovia, Segovia), Juan Pablo Tartari (Fundació Docència i Recerca Mútua de Terrassa and Movement Disorders Unit, Department of Neurology, University Hospital Mutua de Terrassa, Terrassa, Barcelona), Cristina Tejera-Parrado (Instituto de Biomedicina de Sevilla (IBiS), Hospital Universitario Virgen del Rocío/CSIC/Universidad de Sevilla, Seville), Eduard Tolosa (Hospital Clinic Barcelona), Francesc Valldeoriola (Hospital Clinic Barcelona), Laura Vargas-González (Instituto de Biomedicina de Sevilla (IBiS), Hospital Universitario Virgen del Rocío/CSIC/Universidad de Sevilla, Seville), Lydia Vela (Department of Neurology, Hospital Universitario Fundación Alcorcón, Madrid), Francisco Vives (Centro de Investigacion Biomedica, Universidad de Granada, Granada).

**Austria:** Alexander Zimprich (Department of Neurology, Medical University of Vienna, Austria)

**Norway:** Lasse Pihlstrom (Department of Neurology, Oslo University Hospital, Oslo, Norway), Mathias Toft (Department of Neurology and Institute of Clinical Medicine, Oslo University Hospital, Oslo, Norway)

**Estonia**: Pille Taba (Department of Neurology and Neurosurgery, University of Tartu, Tartu, Estonia)

**Australia:** Sulev Koks (Centre for Molecular Medicine and Innovative Therapeutics, Murdoch University, Murdoch, 6150, Perth, Western Australia; The Perron Institute for Neurological and Translational Science, Nedlands, 6009, Perth, Western Australia)

**Israel:** Sharon Hassin-Baer (The Movement Disorders Institute, Department of Neurology and Sagol Neuroscience Center, Chaim Sheba Medical Center, Tel-Hashomer, 5262101, Ramat Gan, Israel, Sackler Faculty of Medicine, Tel Aviv University, Tel Aviv, Israel)

**Finland:** Kari Majamaa (Institute of Clinical Medicine, Department of Neurology, University of Oulu, Oulu, Finland; Department of Neurology and Medical Research Center, Oulu University Hospital, Oulu, Finland), Ari Siitonen (Institute of Clinical Medicine, Department of Neurology, University of Oulu, Oulu, Finland; Department of Neurology and Medical Research Center, Oulu University Hospital, Oulu, Finland), Pentti Tienari (Clinical Neurosciences, Neurology, University of Helsinki, Helsinki, Finland, Helsinki University Hospital, Helsinki, Finland)

**Nigeria:** Njideka U. Okubadejo (University of Lagos, Lagos State, Nigeria), Oluwadamilola O. Ojo (University of Lagos, Lagos State, Nigeria),

**Kazakhstan:** Coordinator - Rauan Kaiyrzhanov (Department of Molecular Neuroscience, UCL Institute of Neurology, London, UK), Chingiz Shashkin (Kazakh National Medical University named after Asfendiyarov, Almaty, Kazakhstan), Nazira Zharkinbekova (South Kazakhstan Medical Academy, Shymkent, Kazakhstan), Vadim Akhmetzhanov (Astana Medical University, Astana Kazakhstan), Akbota Aitkulova (National Center for Biotechnology, Astana, Kazakhstan; Al-Farabi Kazakh national university. Almaty Kazakhstan), Elena Zholdybayeva (National Center for Biotechnology, Astana, Kazakhstan), Zharkyn Zharmukhanov (National Center for Biotechnology, Astana, Kazakhstan), Gulnaz Kaishybayeva (Scientific and practical center “Institute of neurology named after Smagul Kaishibayev”, Almaty, Kazakhstan), Altynay Karimova (Scientific and practical center “Institute of neurology named after Smagul Kaishibayev”, Almaty, Kazakhstan), Talgat Khaibullin (Semey Medical University, Semey, Kazakhstan).

**Ireland:** Timothy L. Lynch (The Dublin Neurological Institute at the Mater Misericordiae University Hospital, Dublin, Ireland & School of Medicine and Medical Science, University College Dublin, Dublin, Ireland).
